# Body mass and latitude predict the presence of multiple stressors in global vertebrate populations

**DOI:** 10.1101/2020.12.17.423192

**Authors:** Nicola Noviello, Louise McRae, Robin Freeman, Chris Clements

## Abstract

Multiple stressors are recognised as a key threat to biodiversity, but our understanding of what might predispose species to multiple stressors remains limited. Here we analyse a global dataset of over 7000 marine, freshwater, and terrestrial vertebrate populations, alongside species-specific trait data, to identify factors which influence the number of stressors a species is subjected to at the population level. We find that body mass and latitude can both influence the number of stressors a population is subjected to across ecosystems, with large-bodied species tending to be more threatened, except terrestrial amphibians which show the opposite trend. Latitudinal forecasts predict higher stressor numbers between 20°N and 40°N, and towards the poles. Global stressor distributions suggest a link between human population centres and stressor frequency generally impacting larger-bodied species. Latitude and body mass hence provide key predictive tools to identify which vertebrate populations are likely to be highly threatened, despite the strength of these trends differing between ecological system and taxonomic class.

## Introduction

Global vertebrate populations are in decline (WWF 2018; He *et al.* 2019), a fact attributed to the overarching trend of anthropogenic stressors (or ‘threats’) increasing in line with human populations (Johnson *et al.* 2017). Such threats can act at both a local and global scale, with locally detrimental stressors such as the overharvesting of species for food (Zhou & Smith 2017) occurring simultaneously against a backdrop of global stressors such as climatic change which can alter energy flows (Bartley *et al.* 2019), impact trophic interactions (Smoliński & Glazaczow 2019), and increase the physiological stress of populations (Iknayan & Beissinger 2018). Thus, as human populations increase, biodiversity is being exposed not only to increasing levels of stress, but to multiple stressors impacting simultaneously; with resulting effects creating novel challenges for the effective conservation of species (Côté *et al.* 2016). Our ability to inform conservation in the face of multiple stressors has been limited by an incomplete understanding of their interactive effects (Darling *et al.* 2013). However, whilst there remains a deficiency in our understanding of exactly how multiple stressors will impact biodiversity, there is mounting concern that stressor interactions will be responsible for increasing disruption to community assemblages (Zavaleta *et al.* 2009). Consequently, a central challenge exists to identify how and why populations are exposed to multiple stressors (Hodgson *et al.* 2017).

Identifying ecological and biotic factors which influence the number of stressors a population is threatened by has clear conservation implications, potentially allowing species to be prioritised – at least for initial appraisal – without the need to collect detailed data at the site or population level. Achieving this requires identifying predictors where data are widely available, and which may *a priori* influence a species’ predisposition to multiple stressors. Here, body mass holds some promise; labelled a ‘supertrait’ (Bribiesca *et al.* 2019) due to its role in numerous ecological processes, wide availability, and universality, it provides a convenient tool enabling direct comparison between taxa. Further linked with an increased vulnerability to strong population decline (Deinet *et al.* 2020) and extinction, body mass has also been suggested as a potential predeterminant of stressor exposure (Collen *et al.* 2011). The reasons for larger bodied organisms being at higher risk of facing one or more stressors are multifaceted, ranging from an increased conspicuousness exposing species to exploitation via human consumption or recreational hunting practices (Verde Arregoitia 2016; Ripple *et al.* 2019); wider home ranges with high resource requirements and subsequent vulnerabilities to habitat degradation (Böhm *et al.* 2016), and higher trophic level increasing vulnerability to cumulative disturbances lower in the food chain (Purvis *et al.* 2000). Stressor exposure may also be affected by ecological system, as marine, terrestrial and freshwater systems each demonstrate their own vulnerabilities to anthropogenic threats. For example, freshwater species may be more commonly subject to nutrient enrichment (Birk *et al.* 2020), whilst exploitation and climate change represent more pressing threats for large species (Halpern *et al.* 2019). Land-use change was shown to be the most common terrestrial threat (Tilman *et al.* 2017) with declines in biodiversity often attributed to the alteration of habitats (Newbold *et al.* 2015).

Neither local nor global stressors are uniformly distributed in space (Bowler *et al.* 2020). Because many local stressors are intimately linked to human populations (e.g., habitat loss, hunting, etc.) the presence and frequency of these stressors is likely to change in line with human population density (Santini *et al.* 2017). The global distribution of human population density varies dramatically between latitudinal belts, with less than 12.5% living below the equator, but around half residing within the comparatively narrow belt between 20°N and 40°N (Kummu & Varis 2011); equating to around 3.85 billion people dwelling within 20 degrees of latitude (United Nations 2019). However, the global effects of climate change are also known to be non-uniform, with much impact expected in the rapid warming of high-latitude, arctic regions (Serreze & Barry 2011), alongside the recent poleward expansion of the tropics and dryer conditions in mid-latitude regions (Staten *et al.* 2018). Consequently, latitude is likely to be a significant determinant of where stressors occur, and thus identifying how stressor number changes on a latitudinal basis would provide further detail on broad threat variability, helping local efforts to mitigate interactions between global and localised stressors.

Disentangling how life history traits, and the spatial heterogeneity of stressors, impact vertebrate populations provides the opportunity to enhance stressor mitigation by combining latitude and body mass, allowing finer differentiation and enabling area-based action across taxa by providing a framework applicable to vertebrates worldwide. Here we take a novel approach in the study of multiple stressors by seeking to identify factors which predict the number of stressors a population is affected by across freshwater, marine, and terrestrial systems at a global scale. To do this we make use of recently available population-level threat data from the Living Planet Database, supplemented with data on body mass from multiple sources, to generate a composite, spatially explicit database for more than 7400 vertebrate populations, comprising 2500 species, across seven continents and all key ecological systems. We use data from the six major taxonomic classes: amphibians, birds, bony fish, cartilaginous fish, mammals, and reptiles, to test whether body mass and latitude alter the number of stressors a population is affected by. We show how latitude and body mass can predict the number of stressors a population is exposed to, with forecasts varying by vertebrate class and ecological system. This research provides new insight on the determinants of multiple stressors in vertebrates, using widely available data on body mass and latitude to provide prioritisation tools for species conservation.

## Materials and Methods

### Data

#### The Living Planet Database

The Living Planet Database (LPD, http://livingplanetindex.org/data_portal) contains information on over 25,000 vertebrate populations around the world, comprising all vertebrate classes across marine, freshwater and terrestrial systems and providing population-specific information such as spatial location, abundance and threat exposure. Data are collected from scientific literature, online databases and grey literature published since 1970, and included if at least two years of abundance records are present, assuming comparable data collection methodologies are used throughout; detailed inclusion criteria for the LPD can be found in Collen *et al.* 2009.

Of the 25,054 population time series making up the LPD (including confidential records), 7826 contained data relating to population threat exposure; comprising up to three of the following stressors: climate change, disease, exploitation, habitat degradation or destruction, habitat loss, invasive species or genes, and pollution. These stressors were counted for each population, with values from a minimum zero, to a maximum of three. Latitude, ecological system, and taxonomic class variables from the LPD were also included in analysis.

#### Body Mass Data

Body mass data were collated from a number of pre-existing databases and the scientific literature (see Supporting Information Table 1 for a full list of sources utilised). Where minimum and maximum values where given, maximum was taken to ensure measures were most likely those of mature individuals, and thus in line with commonly reported measures from the other databases. The majority of data sources did not contain sex-specific body mass measurements; however, where sex was indicated an average of the male / female record was taken to account for dimorphism. Finally, where multiple records of the same species were present between datasets, the mean was taken, with all records then standardised to reflect a common unit (g, grams).

**Table 1.** Model interaction term selection process.

| **Group** | **Models** | **df** | **AICc** | **$\Delta\text{AICc}$**<br>**($\leq 2$)** |
| --- | --- | --- | --- | --- |
| *Terrestrial* |  |  |  |  |
| Amphibians | **NS ~ BS + (1\|S)** | 4 | 261.572 | 0.000 |
| Birds | **NS ~ BS + L + BS:L + (1\|S)** | 12 | 2862.922 | 0.000 |
| Mammals | **NS ~ BS + L + (1\|S)** | 8 | 3593.371 | 0.000 |
| Reptiles | NS ~ BS + L + (1\|S) | 8 | 442.752 | 0.000 |
|  | **NS ~ BS + L + BS:L (1\|S)** | 12 | 443.176 | 0.424 |
|  | NS ~ L + (1\|S) | 7 | 443.744 | 0.992 |
| *Freshwater* |  |  |  |  |
| Amphibians | **NS ~ BS + L + BS:L + (1\|S)** | 12 | 556.038 | 0.000 |
| Birds | NS ~ L + (1\|S) | 8 | 1450.655 | 0.000 |
|  | **NS ~ BS + L + (1\|S)** | 9 | 1452.013 | 1.358 |
| Bony Fish | **NS ~ BS + L + BS:L + (1\|S)** | 13 | 1171.734 | 0.000 |
| Mammals | **NS ~ BS + L + (1\|S)** | 8 | 194.258 | 0.000 |
| Reptiles | NS ~ L + (1\|S) | 8 | 387.802 | 0.000 |
|  | **NS ~ BS + L + (1\|S)** | 9 | 389.466 | 1.663 |
| *Marine* |  |  |  |  |
| Birds | **NS ~ BS + L + BS:L + (1\|S)** | 12 | 1951.925 | 0.000 |
| Bony Fish | **NS ~ BS + L + BS:L + (1\|S)** | 12 | 2135.723 | 0.000 |
| Cartilaginous Fish | **NS ~ BS + L + (1\|S)** | 8 | 282.502 | 0.000 |
| Mammals | No parameters suggested | 3 | 771.875 | 0.000 |
|  | **NS ~ BS + (1\|S)** | 4 | 773.391 | 1.516 |
|  | NS ~ L + (1\|S) | 7 | 773.793 | 1.918 |
| Reptiles | **NS ~ L + (1\|S)** | 8 | 492.981 | 0.000 |

Model selection process to determine appropriate interactive terms. Models were ranked according to their corrected Akaike Information Criterion (AICc) values, with models $\leq \Delta\text{AICc } 2$ shown for each class and ecological system. NS = number of stressors; BS = body size (ln of mass in grams); C = taxonomic class; L = latitude; S = species; colons (:) represent interactive terms. Models selected for use in analysis are highlighted in bold.

#### Body Mass Estimations

For some taxa body mass data were unavailable, and so were estimated using allometric regression equations with clade and measurement□specific values, where possible (Feldman *et al.* 2016; Stark *et al.* 2020). These were configured *W = a L^b^*, where *W* = body mass, *L* = length and priors *a* and *b* are the intercept and slope of a regression line over log □ transformed weight □ at □ length data, respectively (Ripple *et al.* 2017; Froese & Pauly 2019). This method was applied to 47 amphibian species using snout to vent length (SVL) records and clade-specific priors (Santini *et al.* 2018; Stark *et al.* 2020), and a further 320 fish species’ mass were estimated, based on maximum total length (TL) and regression priors, as listed on FishBase (Froese & Pauly 2019). Where a measure other than TL was listed (e.g., standard length (SL), fork length (FL)), regression coefficients were used to convert these to total length before then estimating body mass.

#### Final Dataset

Upon the merging of body mass and LPD threat data, the final database totalled 7470 population records, representing 2516 vertebrate species; ~ 3.5% of described vertebrate species (IUCN 2020). To normalize residuals and reduce heteroscedasticity, body mass (g) data were log transformed prior to the analyses (Baliga *et al.* 2019; Stark *et al.* 2020). The full dataset was partitioned by taxonomic class and ecological system, providing 14 subsets used for model building, with each representing one group e.g., marine birds, terrestrial mammals, etc. Freshwater cartilaginous fish were discounted from this procedure due to a paucity of data.

### Statistical Analysis

We explored a model relating the number of threats to body mass and latitude and their interactions *(number of threats ~ body mass * latitude).* We used a generalised linear mixed modelling (GLMM) framework with a truncated Conway-Maxwell Poisson family using the ‘glmmTMB’ package (Brooks *et al.* 2017). This accounted for the discrete, count data of stressor number present for each population. Our model building strategy employed the use of existing knowledge (e.g., Ripple *et al.* 2017; Bowler *et al.* 2020) to construct a logical and plausible set of *a priori* predictors to describe relationships between stressor number and traits which predispose populations to vulnerability of stressors.

Prior to analysis, latitude and the natural log of body mass were evaluated using Variance Inflation Factor (VIF) and were considered beneath the threshold to constitute collinearity in all cases (threshold = 2). The combination of generalized linear models with splined data has been shown to provide a straightforward parametric approach to modelling non-linear terms, whilst performing better than their additive counterparts (He *et al.* 2006; Chung *et al.* 2009). Consequently, natural (restricted) cubic splines with four degrees of freedom, corresponding to five knots (Stone 1986; Shepherd & Rebeiro 2017) were applied to the latitude variable within all relevant models to account for the non-linear relationship with stressor number (see Fig. 1). Multiple intra-specific populations were accounted for using species as random effect, maintaining random intercepts.

**Fig. 1.**
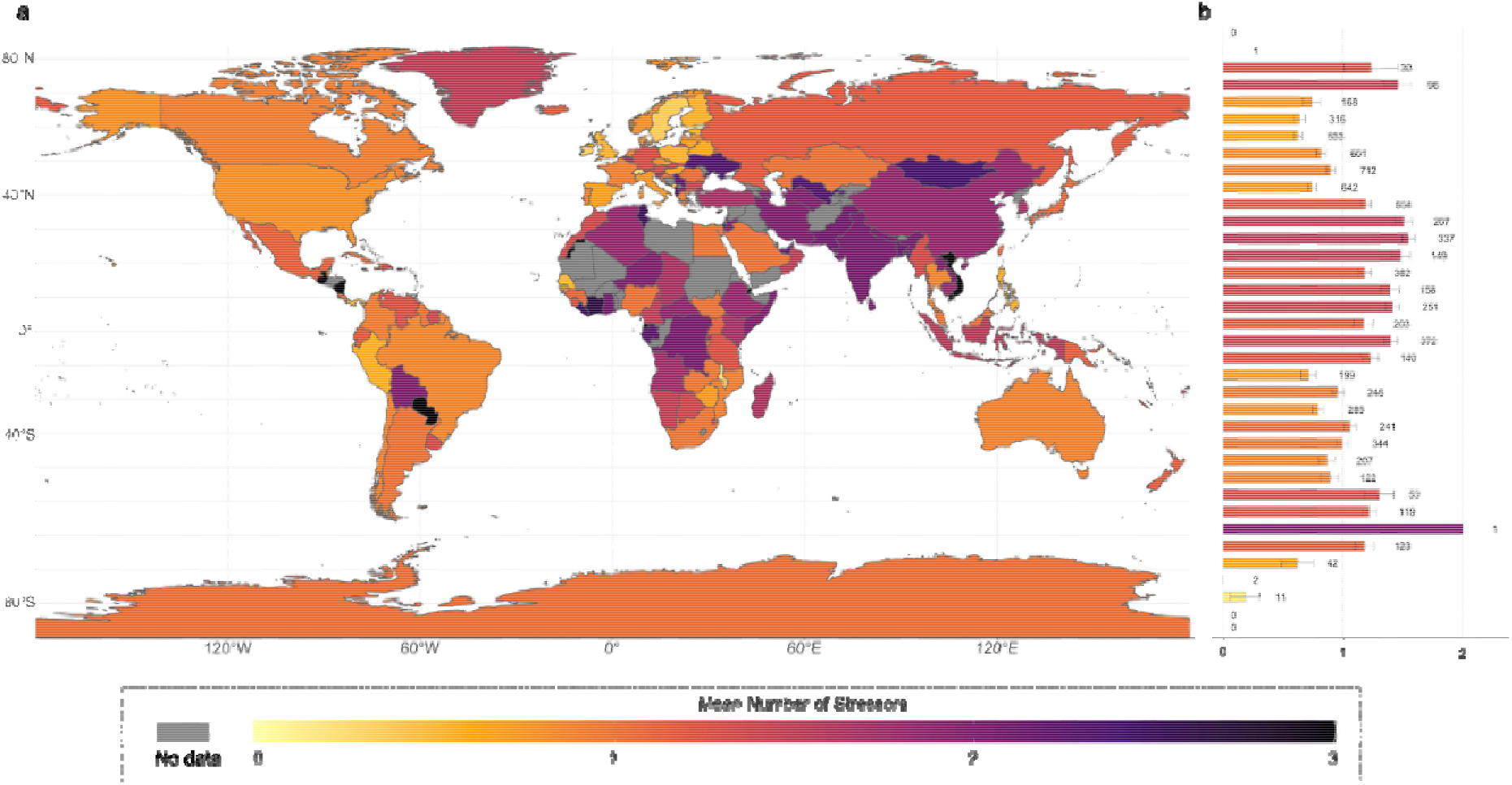
Global distribution of mean number of stressors by country and latitude. Global overview of the mean number of stressors **a** within each country and **b** by latitude with numbers alongside bars representing sample sizes for each 5° latitude bin.

Full models generally failed to converge, and so to identify the best combination of two-way interaction terms for each group, ‘dredge’ from the ‘MuMIn’ package was implemented (Bartón 2014) using the model structure: *number of threats ~ body mass * latitude* and including two-way interactions between body mass and latitude. With corrected Akaike Information Criterion for small sample sizes (AICc) used to identify the best model for each system (Anderson & Anderson 2002), we highlight candidate models with ΔAICc ≤2, and selected the best model for each system scenario, based on AICc and including at least one predictor where reasonable (see Table 1).

Whilst AICc was used to aid model selection by making inferences from multiple models, at the same time considering fit and complexity (Johnson & Omland 2004), estimates of pseudo-R^2^ were used to assess model fit in the final models using the “r2_nakagawa” function from the “performance” package (Nakagawa & Schielzeth 2013) (Table 2). All statistics were performed in R V3.6.1.

**Table 2.**
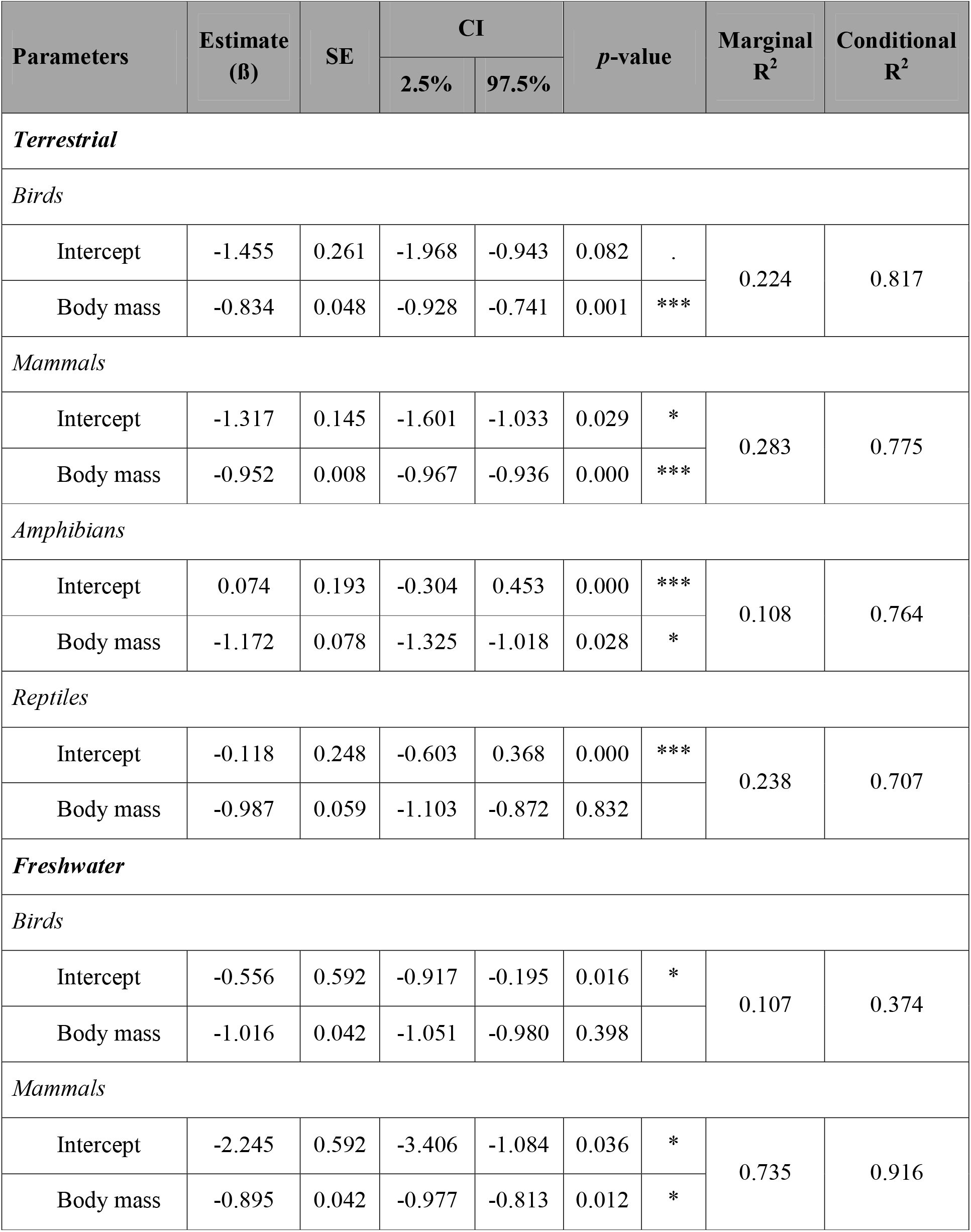
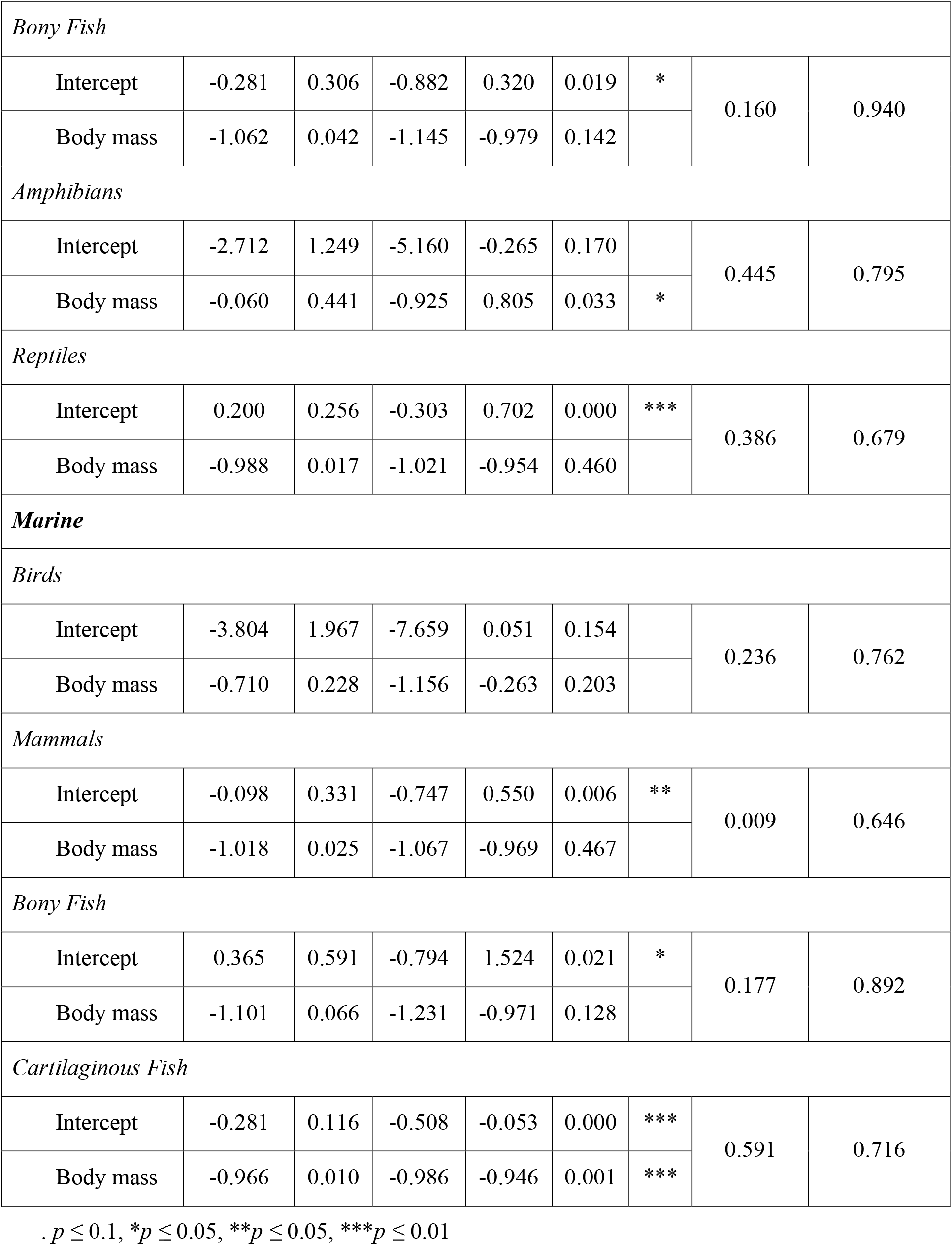
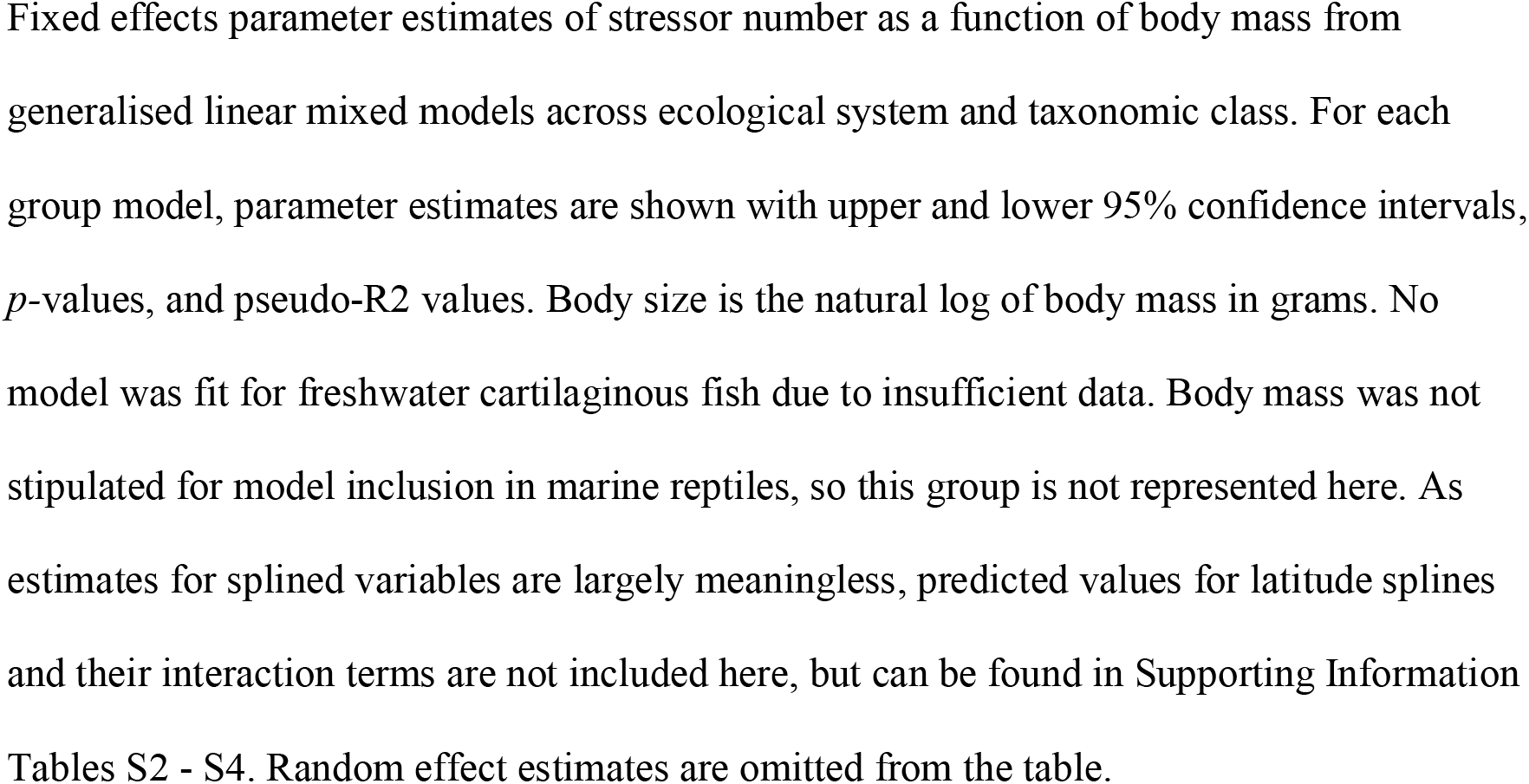
Summary of generalised linear mixed model coefficients for body mass parameter for each class and ecological system.

## Results

Stressor number showed a clear latitudinal gradient, with midrange latitude populations typically subject to fewer stressors than those between 10°S and 30°N, or at the poles (Fig. 1). Countries with highest mean number of stressors are typically those with high human population density and large population sizes, notably China and the Indian subcontinent. Indeed, of those populations affected by three stressors, most (23.53%) were situated in Asia which also showed the highest mean number of stressors across populations (1.53). Conversely, the highest proportion of populations subject to zero stressors (31.54%) were situated in Europe which also had the lowest mean number of stressors per population (0.74). The number of stressors affecting a population varied with body mass, with smaller species typically having fewer threats (Fig. 2). The noticeable exception to this was in amphibians, where the heaviest species were generally affected by fewer stressors (59 populations; Fig. 2b). Single stressors were observed in 35.0% of vertebrate populations, but most frequently seen in mid-sized birds, mammals, reptiles and fish species. Again, amphibians differed from this pattern, showing a more normal body mass distribution across populations and a reduced exposure to single stressors when viewed alongside other vertebrate classes (Fig. 2b).

**Fig. 2.**
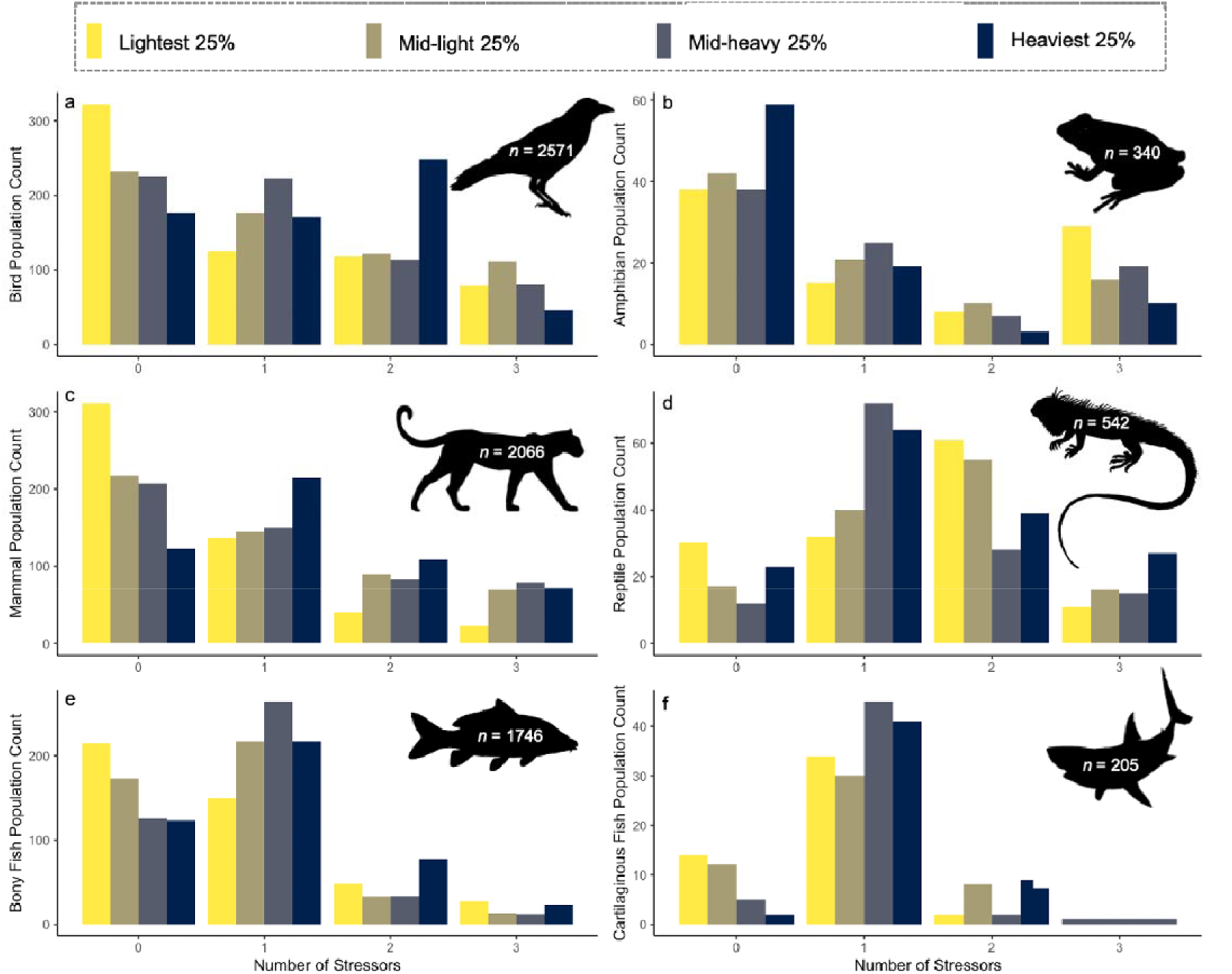
Number of stressors by body mass quartile. Populations exposed to each number of stressors (0 - 3) shown as counts for the lightest 25%, mid-light 25%, mid-heavy 25% and heaviest 25% species, within each taxonomic class. Panels represent **a** birds **b** amphibians **c** mammals **d** reptiles **e** bony fish and **f** cartilaginous fish.

Populations exposed to two stressors accounted for 18.0% of records with the heaviest bird, mammal and bony fish populations (249, 109 and 78, respectively) more likely to be impacted than those with a lower body mass, with bird species demonstrating the clearest distinction (Fig. 2a). Exposure to two stressors differed in amphibians and cartilaginous fish, while 33% of reptile populations were exposed to two stressors; more than any other group. Populations were least likely to be exposed to three stressors (10.3% overall), with fish having the fewest populations subjected to three stressors (bony fish 77 populations; cartilaginous fish just one; Fig. 2e and f). Remaining classes showed similar exposure levels but differed throughout body mass ranges, with no clear pattern demonstrated.

Consistent with patterns demonstrated by body mass quartiles (Fig. 2), our models produced several significant positive relationships between species body mass and the likelihood of populations being exposed to higher stressor numbers, except in one group for which fewer stressors were predicted as body mass increased (Table 2 and Fig. 3). Terrestrial systems showed some of the clearest relationships (Fig. 3a), with predicted stressor number significantly increasing with body mass for terrestrial mammals and birds (ß□=□ −0.952 ± 0.008, *p*-value □ = <0.001 and ß□=□ −0.834 ± 0.048, *p*-value = 0.001, respectively). Meanwhile, terrestrial amphibians demonstrated a negative relationship between stressor number and body mass (ß□=□-1.172 ± 0.078, *p*-value □ = 0.028). Cartilaginous fish produced the only significant estimates in marine ecosystems (ß□= −0.966 ± 0.010, *p*-value□=□0.001, Fig. 3b) while freshwater mammals and amphibians showed similar trends, with predictions forecasting more stressors as body mass increases (ß□=□-0.895 ± 0.042, *p*-value = 0.012 and ß□=□ −0.060 ± 0.441, *p*-value□ = 0.033, respectively. Fig 2.3c.).

**Fig. 3.**
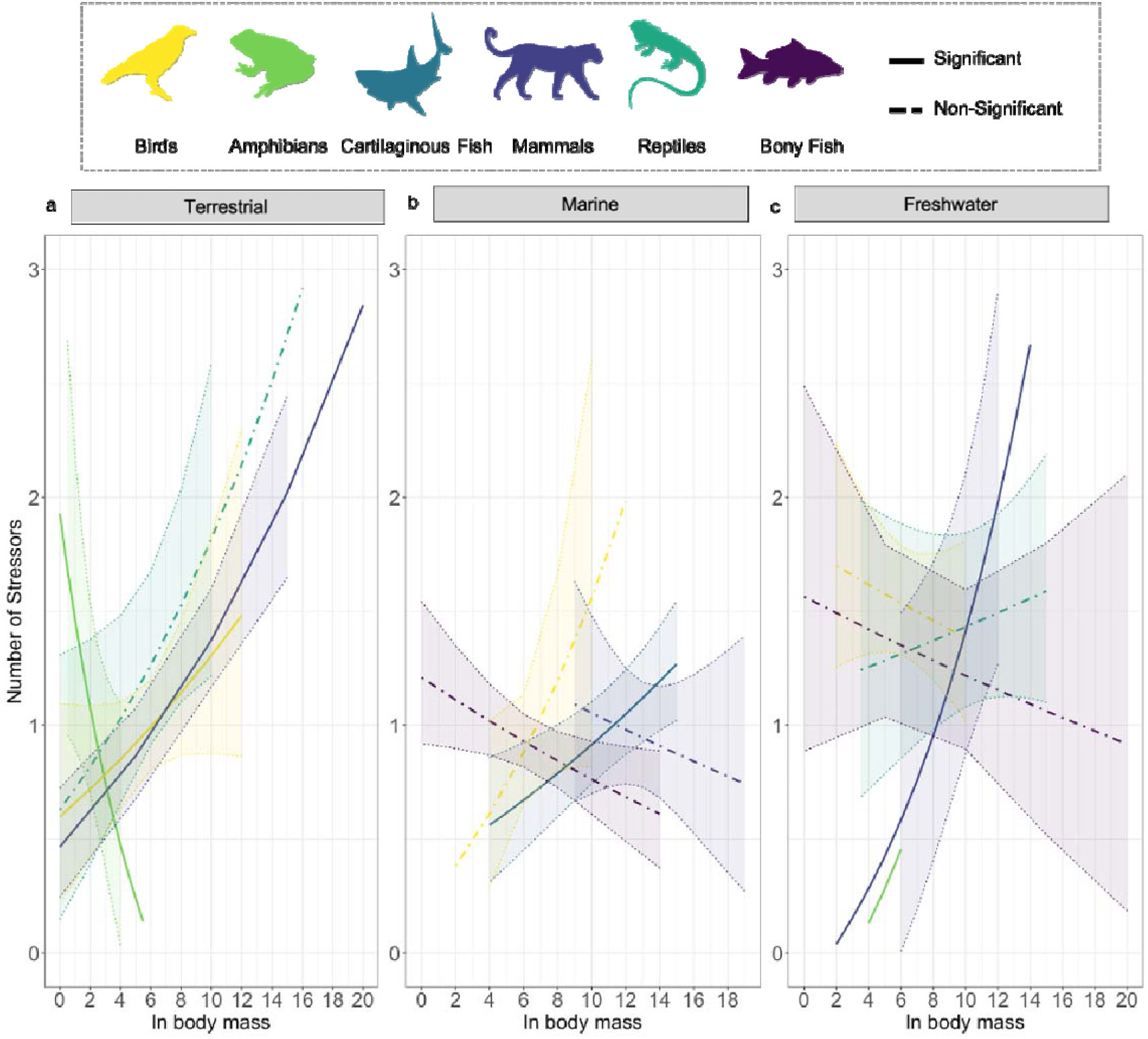
Model predictions of stressor number as a function of body mass. Generalized linear mixed model predictions of stressor number as a function of natural log of body mass in grams for **a** terrestrial ecosystems **b** marine ecosystem, and **c** freshwater ecosystems. Class predictions with *p* = ≤0.05 are represented by a solid line whilst non-significant estimates are shown by dashed lines. Ribbons display 95% confidence intervals.

Model predictions of stressor number as a function of latitude showed varied results, but with some common patterns (Fig. 4). For the majority of groups (e.g., marine mammals, terrestrial reptiles, freshwater amphibians) the most stressors were experienced towards extreme latitudes. This pattern is often reversed at the opposite pole, with the fewest stressors predicted. For instance, marine bony fish experience the most stressors towards the southern pole, but fewest in northernmost latitudes (Fig. 4b), a trend which is reversed in freshwater mammals, where more stressors are predicted at northern latitudes, and fewer towards southern extremities. There were also several groups, across systems, with local maxima between approximately 20°N and 40°N, such as terrestrial mammals and reptiles, marine cartilaginous fish and freshwater birds, mammals, amphibians and reptiles.

**Fig. 4.**
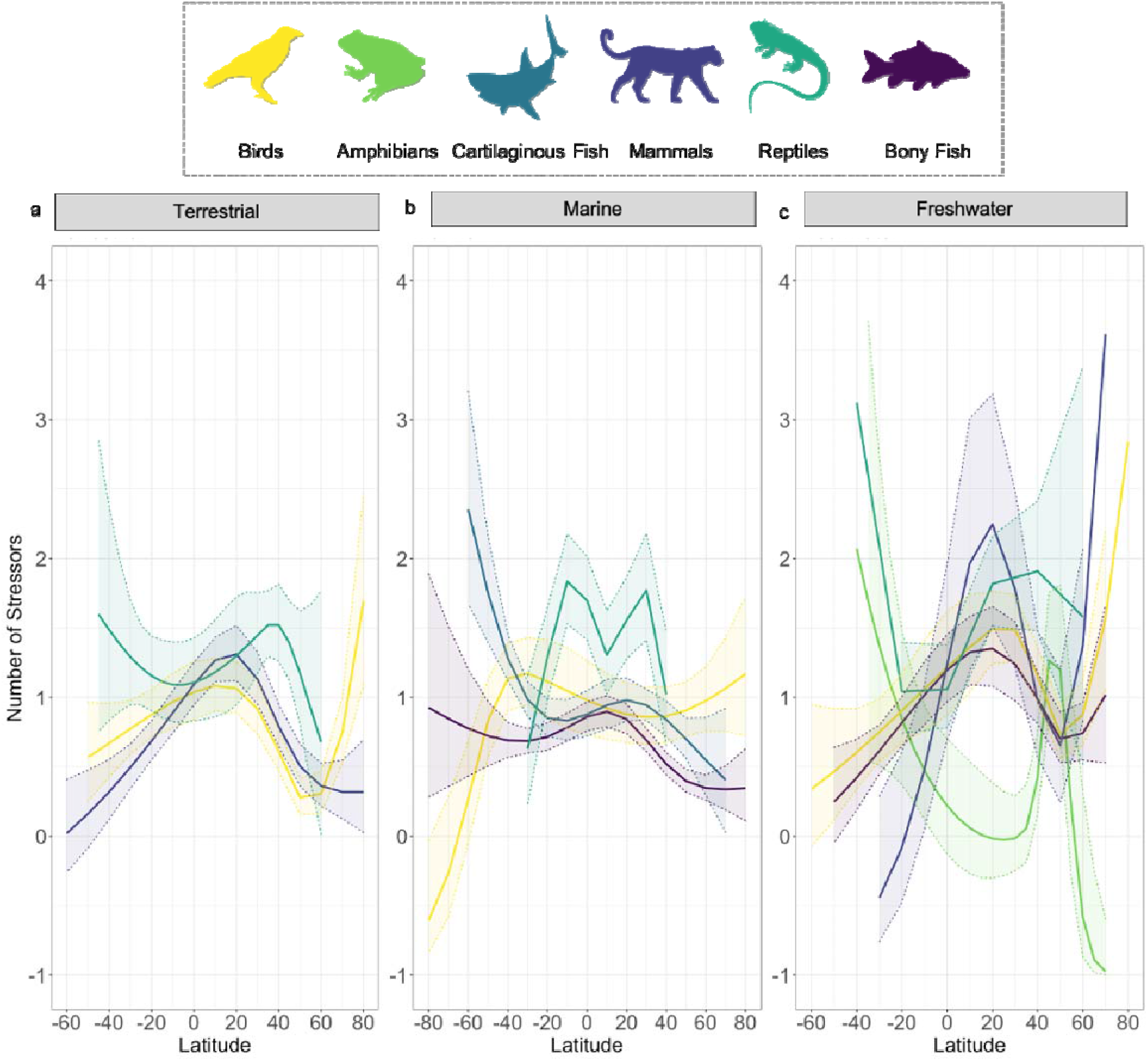
Model predictions of stressor number as a function of latitude. Generalized linear mixed model predictions of stressor number as a function of natural log of body mass in grams for **a** terrestrial ecosystems **b** marine ecosystem, and **c** freshwater ecosystems. Solid lines represent parameter estimates, with ribbons showing 95% confidence intervals. Splined parameter predictions cannot be numerically interpreted, and so all estimates are displayed graphically here for interpretative purposes.

Stressor number also varied considerably by latitude in interaction with body mass (Fig. 5). Again, peaks and lows were predicted towards polar regions for marine birds (Fig. 5b), and freshwater amphibians (Fig. 5c). However, interactions between latitude and body mass also seem to play a part in stressor exposure for most classes, with the heaviest quartile tending to experience the most stressors regardless of latitude, except in amphibians, where it is the lightest quartile that are predicted to suffer higher stressor numbers, except at lower latitudes where the heaviest seem most at risk of stressor exposure (Fig. 5c). Freshwater bony fish suffer greatest stressors between 10°S and 30°N, with local peaks apparently dependent on body mass, as larger species suffer greater stressors at lower latitudes and lighter taxa reaching peaks at around 20°N, and again towards northern extremities (Fig. 5c). Heavier marine birds also suffer a peak in stressor number at 40°S, while the first quartile peaks at around 30°N (Fig. 5b). Marine bony fish show a gentle summit at around 10°N, but body mass seems to have a lesser effect than in other classes (Fig. 5b).

**Fig. 5.**
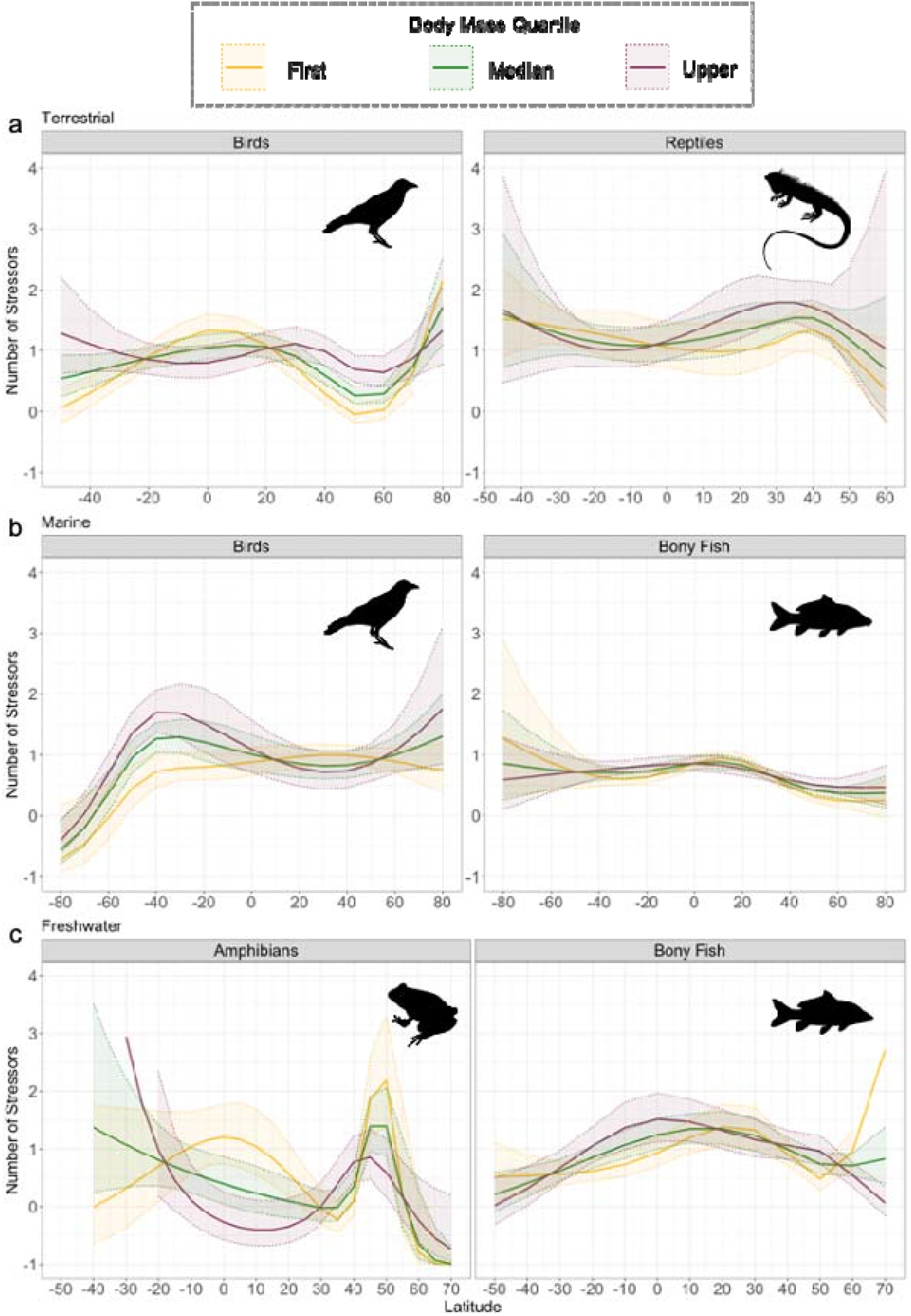
Model predictions of stressor number as a function of latitude by interaction with body mass. Generalized linear mixed model predictions of stressor number as a function of latitude with splines and four degrees of freedom for **a** terrestrial birds and reptiles **b** marine birds and bony fish, and **c** freshwater amphibians and bony fish. Solid lines represent parameter estimates, with ribbons showing 95% confidence intervals. Splined parameter predictions cannot be numerically interpreted, and so all estimates are displayed graphically here for interpretative purposes.

While plots are provided, note that coefficient estimates for the latitude variable and its interactive effects are absent from Table 2. As interpretation of splined variable coefficients is essentially meaningless (Shepherd & Rebeiro 2017) our predictions are instead displayed visually in Fig. 4 and Fig. 5 to elucidate our findings. Full coefficient estimates and intercept values have been provided within Supplementary Tables 2, 3 and 4.

## Discussion

An inability to identify correlates of exposure to multiple stressors could hinder broad efforts to maintain biodiversity and the mitigative actions implemented to protect species (Hodgson *et al.* 2017). Using a new and comprehensive collation of vertebrate body mass data, alongside population records taken from the Living Planet Database (LPD), we show that both body mass and latitude can provide valuable estimates of stressor number in global vertebrate populations. Our results reveal class and system specific patterns in stressor number as a function of body mass, and latitude. Spatial patterns show mean stressor number is highest between 20°N and 40°N, and again towards the poles, with model predictions further supporting these latitudinal findings while producing more nuanced predictions of stressor number when combined with body mass differentiated by ecological system and class. Uncovering the relationship between body mass and number of threats has useful applications for vertebrate conservation. Though we know comparatively little about the world’s rarest species because of their scarcity (Ripple *et al.* 2017), an easily measured and universal predictor such as body mass, provides an accessible tool for approximating threat status where little other data exists. Likewise, latitude provides a secondary measure and greater specificity to population threat estimates. As such, we provide an adaptive framework for further exploration of stressor predictors, using ubiquitous traits both biological and geographical.

Our models revealed positive relationships between body mass and the number of stressors affecting terrestrial mammals and birds, marine cartilaginous fish, freshwater mammals and amphibians within the LPD. This trend was generally anticipated, given the well-documented challenges faced by large-bodied species. These include heightened levels of exploitation, damage to habitats and larger home range increase the chances of stressor exposure in heavier species (Collen *et al.* 2011; Böhm *et al.* 2016). This is further supported by complementary research from the Living Planet Index, which most recently demonstrated a link between larger body mass and strong population declines in freshwater ecosystems(He *et al.* 2019). Conversely, terrestrial amphibians showed a significant negative relationship where lower body mass predicted higher stressor number (Fig. 3), a pattern further supported by raw data shown in Fig. 2b. Prior research has already produced instances of amphibians standing alone in threat trends when compared to other vertebrate classes, with the lightest amphibian species subject to higher instances of threats signifying habitat degradation or loss, than heavier species (Ripple *et al.* 2017). Smaller terrestrial species may also generally have a limited dispersal ability, restricted geographic range (Cardillo *et al.* 2008) and occupy narrow niches incapable of enduring change (González-Suárez & Revilla 2013). Arboreal amphibians are also the lightest of their class (Santini *et al.* 2018) and are likely more prone to impacts from deforestation and land clearing. It remains uncertain why amphibians are the apparent exception amongst vertebrates with regards exposure to multiple stressors, but more data would enable the specialist analysis required to further explain whether amphibian declines are due to the currently suspected fungal disease (Carvalho *et al.* 2017), or disease in combination with additional stressors.

Previous work has highlighted that predators are particularly at risk of anthropogenic change (Ripple *et al.* 2017), with the effect that predators are being disproportionately lost worldwide (Estes *et al.* 2011). Given that body mass is often positively related with trophic level, particularly in aquatic environments (Riede *et al.* 2011), our analysis suggest that predators are likely to be disproportionately threatened by multiple stressors, which may in part account for these observed losses. These impacts on predators are known to have cascading effects on the stability of food webs by changing the strength of direct and indirect top-down effects (McCary *et al.* 2020). By altering the stability of predator populations, changes in biodiversity (Levin & Lubchenco 2008), biomass (Soliveres *et al.* 2016), disease and carbon sequestration are all possible (Estes *et al.* 2011), with extensive cascading effects seen throughout ecosystems worldwide (Beschta & Ripple 2020). Moreover, with trophic downgrading shown to interact with pre-existing anthropogenic threats such as pollution and habitat change (Estes *et al.* 2011), and pressures becoming ever more prevalent globally (Halpern *et al.* 2015), further integration of trophic level into predictive estimates of stressor number would provide greater insight into these relationships. Nevertheless, with recent work highlighting an increased risk of extinction in some herbivorous groups (Atwood *et al.* 2020), it should be noted that species at the bottom of the consumer triangle can also be powerful influencers within their communities, commanding architectural power over ecosystem structures via the physical removal of vegetation and increased influence on the biomass cycle (Barnosky *et al.* 2016).

Human population density has long been considered a proxy for anthropogenic disturbance factors (Santini *et al.* 2017), with local population driven stressors such as pollution and exploitation leading to an increase in the frequency of stressors in the northern hemisphere where most economic activity takes place (Moore 2016). Median body mass is also generally higher for terrestrial mammals in the northern hemisphere (Santini *et al.* 2017), and with the additional impact of human pressures, heavier species are at a greater probability of being afflicted by multiple stressors. Findings from our latitudinal models (Fig. 4) and Fig. 1 support this throughout, indicating locally increased risks across the northern hemisphere where human population is densest. Indeed, some of the world’s densest cities – including Mumbai, Taipei, Shanghai, Karachi and Dhaka – fall between 19°N and 34°N, with around 50% of the human population living between 20°N and 40°N (Kummu & Varis 2011). Notable exceptions in this pattern are marine birds and, arguably, marine reptiles which show a bimodal latitudinal pattern; although the latitudinal gradient in the latter may not be of sufficient span to provide an unequivocal pattern. For marine birds, it is plausible that many species are pelagic and so avoid coastal areas associated with the highest levels of population density and greatest cumulative impact of human activity (Halpern *et al.* 2008). With fishing pressures greatest in the northern hemisphere (Kroodsma *et al.* 2018), and fewer people below the equator (Kummu & Varis 2011) climate-related threats may be more influential than suspected in some groups. This is demonstrated in populations showing southerly maxima in predicted stressor number (such as marine mammals, and terrestrial reptiles), as minimal human population density infers scant local stressors in the southern hemisphere. Indeed, several classes were predicted an increase in stressor number further towards at least one polar region (see terrestrial reptiles and birds, marine mammals, and freshwater reptiles, birds, amphibians and mammals; Fig. 4a-c). As estimates differ by class and by body mass quartile, this may indicate differing abilities of taxa in coping with climate-related changes to niches, with polar amplification a pertinent concern for many species whose ranges reach into Arctic and Antarctic regions (Vincent 2019). Species reliant on sea ice, the Pacific walrus *(Odobenus rosmarus)* for instance, might be subject to both climate change pressures and degradation of habitat as sea ice levels experience record minima (Post *et al.* 2013).

Congruent with peaks in mean stressor number (Fig. 1) and latitudinal stressor predictions (Fig. 4) are several low-and middle-income economies, with a suite of socio-economic issues leading to heightened stressor numbers and a compromised ability to mitigate threats (Vörösmarty *et al.* 2010). For instance, rapidly developing countries typically see surges in infrastructure, pollution, poor regulation of exploitative activities such as hunting, and further unsustainable use of natural resources, which can all increase the stress exerted on biological systems (Vörösmarty *et al.* 2010; Venter *et al.* 2016). The lack of regulatory guidance on hunting for food and medicinal products of particular concern, and with nearly all threatened mammals occurring in developing countries (Ripple *et al.* 2016), large threatened species are again more likely to be impacted than lighter less endangered groups. Moreover, wealthy countries commonly export their waste processing abroad, effectively outsourcing their environmental impact to lower income countries which are less able to deal with the effect (Brooks *et al.* 2018). Yet it is also plausible that threatened species are over-represented in low- and middle-income countries, where a paucity of conservation funding compels prioritisation towards species considered most at-risk. Despite more developed countries demonstrating generally lower mean stressor number in this study, existing research suggests that even high stressor numbers tend to be tolerated until their impacts become detrimental, at which point it is generally not the cause of stressors treated, but the symptoms (Vörösmarty *et al.* 2010). Alas, this pattern may be more consequential for developing countries which will likely have to manage the repercussions, thus further increasing already high stressor numbers for low- and middle-income nations.

Human activity is widely considered liable for the incipient sixth mass extinction (Ceballos *et al.* 2020). The long history combining highly modified landscapes alongside exploitation, climate change and the introduction of invasive species has already driven many modern species to extinction (Otto 2018), and multiple stressor combinations are forecast to push yet more populations beyond the point of recovery (Symes *et al.* 2018), with large bodied species particularly prone to loss (Otto 2018). As population growth is forecast to undermine the protection of natural environments (Crist *et al.* 2017), humanity presents itself with daunting challenges for multiple stressor mitigation, and resource provision for a quickly expanding populace. Corresponding growth in food demand will likely, therefore, place even greater pressure on the species targeted for harvesting; highlighting not only the plight of favoured, large-bodied vertebrates, but jeopardising the viability of long relied-upon resources should these species be exploited beyond recovery (Ripple *et al.* 2017). Yet, it is suspected that measures developed to curtail such impacts would not be sufficient, with the reduction of food waste, changes to diet, restrictions to the harvesting of wild species, and the intensification (rather that expansion) of food production worthy, but deficient, approaches (Crist *et al.* 2017). The minimisation, and eventual reversal, of population growth has been suggested as one of the few measures capable of generating the changes required to sufficiently manage stressors, whilst maintaining biodiversity and human resource requirements concurrently (Crist *et al.* 2017). Recent research has provided some hope for this strategy, with a fall in human fertility projected to dramatically reduce population growth worldwide (Vollset *et al.* 2020). Notwithstanding the social implications of this, such news may bring some hope for conservationists conscious of the cumulative impact that multiple anthropogenic stressors have had on the natural world since the beginning of the Anthropocene.

This research describes stressor number as a function of body mass for thousands of vertebrate populations, yet it should be noted that analyses were only possible for those with available stressor data for respective populations within the LPD; as such we were able to generate predictions for around 3.5% of described vertebrate species. Thus, despite differentiation by class and system providing some generalisability within specific groups and enabling conservation planning within those groups, caution should be exercised when applying our findings to the management of species beyond our analyses; particularly in classes represented by smaller sample sizes within the LPD. While the LPD draws from published literature, this also means its data inherits any biases derived from its sources. This has resulted in the over-representation of well-studied regions and taxa, with research also inclined towards populations within protected areas and terrestrial ecosystems (McRae *et al.* 2017). To this end, we advocate explicit reference to threats within ecological research to enable the expansion of current databases and to keep multiple stressor processes at the forefront of developing research; particularly in underrepresented areas classes such as cartilaginous fish and amphibians.

## Conclusion

Body mass and latitude predict the frequency of stressors faced by global vertebrate populations. Our findings that body mass and latitude can be used as predictors of stressor number in vertebrates offer the ability to streamline conservation prioritisation, whilst emphasising opportunities for the dramatic change required to minimise stressors for the future of our planet. With threat level differing by class and ecological system, we present a framework universally applicable, yet capable of distinguishing between multiple species and population-specific factors. Our results support previous research suggesting that the large charismatic creatures fronting conservation initiatives globally are likely to be the species most at risk. Yet by highlighting the plight of large-bodied species, we hope that protection of these ecosystem architects is enhanced and expedited.

## Supporting information

Body Mass Data

## Supporting Information

Supporting Information is available for this paper:

**Table S1 | Body Mass / Length Sources**

**Table S2 | Terrestrial model coefficients**

**Table S3 | Marine model coefficients**

**Table S4 | Freshwater model coefficients**

**Data file S1 | Body mass data**

## Funding

L.M. is funded by WWF UK and WWF Netherlands, C.C. is supported by grant RPG- 2019-368.

## Author contributions

C.C., L.M. and R.F. conceptualised the project. R.F. and L.M curate and maintain the Living Planet Database. N.N. collated body mass data, performed analyses and produced the manuscript. All co-authors contributed substantially to revisions.

## Data and materials availability

The Living Planet Database (excluding confidential records) is available at: www.livingplanetindex.org/data_portal. All other data needed to evaluate the conclusions in the paper are present in the paper and/or the Supporting Information. Code used in this study is available at: www.github.com/nicolanoviello87/LPD.

## Notes

https://livingplanetindex.org/data_portal

https://github.com/nicolanoviello87/LPD

### Competing Interest Statement

The authors have declared no competing interest.

