## Supporting Information for "Body mass and latitude predict the presence of multiple stressors in global vertebrate populations"

**Table S1 | Body Mass / Length Sources**

This table includes all sources of body mass data used within the study, plus measurements used in allometric mass / length equation calculations.

| **Data** | **Measurement** | **Reference** |
| --- | --- | --- |
| **Amniote** | Body Mass | Nathan P. Myhrvold, Elita Baldridge, Benjamin Chan, Dhileep Sivam, Daniel L. Freeman, S. K. Morgan Ernest. 2015. An amniote life-history database to perform comparative analyses with birds, mammals, and reptiles. Ecology 96: 3109 |
| **AmphiBIO** | Body Mass | Oliveira, B., São-Pedro, V., Santos-Barrera, G. et al. AmphiBIO, a global database for amphibian ecological traits. Sci Data 4, 170123 (2017) |
| ***Atelopus longirostris*** | Body mass | Elicio Eladio Tapia, Luis Aurelio Coloma, Gustavo Pazmiño-Otamendi & Nicolás Peñafiel (2017) Rediscovery of the nearly extinct longnose harlequin frog Atelopus longirostris (Bufonidae) in Junín, Imbabura, Ecuador, Neotropical Biodiversity,  3: 1, 157-167, DOI:  10.1080/23766808.2017.1327000 |
| ***Chalcorana (Rana) chalconota*** | SVL | Robert F. Inger, Bryan L. Stuart, Djoko T. Iskandar, Systematics of a widespread Southeast Asian frog, Rana chalconota (Amphibia: Anura: Ranidae), Zoological Journal of the Linnean Society, Volume 155, Issue 1, January 2009, Pages 123–147, https: //doi.org/10.1111/j.1096-3642.2008.00440.x |
| **Elton Traits** | Body Mass | Smith et al 2003, Dunning 2007 – see Elton traits metadata |
| **Encyclopedia of Life** | Body Mass | Parr, C. S., N. Wilson, P. Leary, K. S. Schulz, K. Lans, L. Walley, J. A. Hammock, A. Goddard, J. Rice, M. Studer, J. T. G. Holmes, and R. J. Corrigan, Jr. 2014. The Encyclopedia of Life v2: Providing Global Access to Knowledge About Life on Earth. Biodiversity Data Journal 2: e1079, doi:10.3897/BDJ.2.e1079 |
| **Fishbase** | Length (TL / FL / SL) | Froese R. & Pauly D. (eds). (2020). FishBase (version Feb 2018). In: Species 2000 & ITIS Catalogue of Life, 2020-09-01 Beta (Roskov Y.; Ower G.; Orrell T.; Nicolson D.; Bailly N.; Kirk P.M.; Bourgoin T.; DeWalt R.E.; Decock W.; Nieukerken E. van; Penev L.; eds.). Digital resource at www.catalogueoflife.org/col. Species 2000: Naturalis, Leiden, the Netherlands. ISSN 2405-8858. |
| **Handbook of the Birds of the World Alive** | Body Mass | S. M. Billerman, B. K. Keeney, P. G. Rodewald, and T. S. Schulenberg (Editors) (2020). Birds of the World. Cornell Laboratory of Ornithology, Ithaca, NY, USA. https://birdsoftheworld.org/bow/home |
| ***Leiopelma archeyi*** | Body mass | Stark, G,  Meiri, S.  Cold and dark captivity: Drivers of amphibian longevity. Global Ecol Biogeogr.   2018; 27:  1384– 1397. https: //doi.org/10.1111/geb.12804 |
| ***Litoria australis* aka *Ranoidea australis, Litoria dahlia* aka *Ranoidea dahlii, Ranoidea genimaculata* aka *Litoria genimaculata*** | SVL | Vanderduys, E. (2019). Field Guide to the Frogs of Queensland. In Field Guide to the Frogs of Queensland. doi: 10.1071/9780643108790 |
| ***Litoria nannotis* aka *Ranoidea nannotis*** | Body mass | Liem, D.S. (1974). A review of the Litoria nannotis species group and a description of a new species of Litoria from north-east Queensland, Australia. Memoirs of the Queensland Museum 17(1), 151-168.  Cogger, H.G. (1994). Reptiles and Amphibians of Australia. Reed Books, Sydney.  McDonald, K.R. & Alford, R.A. (1999). A Review of Declining Frogs in Northern Queensland. Pp 14-22 in A. Campbell (ed), Declines and Disappearances of Australian Frogs. Environment Australia, Canberra. 234 pp. |
| ***Myotis escalerai*** | Body mass | Quetglas, J. (2016). Murciélago ratonero ibérico – Myotis escalerai. En:  Enciclopedia Virtual de los Vertebrados Españoles. Salvador, A., Barja, I. (Eds.). Museo Nacional de Ciencias Naturales, Madrid. |
| **PanTHERIA** | Body Mass | Kate E. Jones, Jon Bielby, Marcel Cardillo, Susanne A. Fritz, Justin O'Dell, C. David L. Orme, Kamran Safi, Wes Sechrest, Elizabeth H. Boakes, Chris Carbone, Christina Connolly, Michael J. Cutts, Janine K. Foster, Richard Grenyer, Michael Habib, Christopher A. Plaster, Samantha A. Price, Elizabeth A. Rigby, Janna Rist, Amber Teacher, Olaf R. P. Bininda-Emonds, John L. Gittleman, Georgina M. Mace, and Andy Purvis. 2009. PanTHERIA: a species-level database of life history, ecology, and geography of extant and recently extinct mammals. Ecology 90: 2648. |
| ***Rana tavasensis*** | SVL | Düşen, S. (2012). First data on the helminth fauna of a locally distributed mountain frog, “Tavas frog” Rana tavasensis Baran & Atatür, 1986 (Anura: Ranidae), from the inner-west Anatolian region of Turkey. Turkish Journal of Zoology, 36, 496-502. |
| ***Trachycephalus venulosus*** | Body mass | Domingos J. Rodrigues, Masao Uetanabaro & Frederico S. Lopes (2005) Reproductive patterns of Trachycephalus  venulosus (Laurenti, 1768) and Scinax fuscovarius (Lutz, 1925) from the Cerrado, Central Brazil, Journal of Natural  History, 39: 35, 3217-3226, DOI:  10.1080/00222930500312244 |
| **Various amphibian** | Body Mass | Santini L., Benítez-López A., Ficetola G.F., Huijbregts M.A.J. 2017. Length – Mass allometries in Amphibians. Integrative Zoology, 13: 36-45. doi:10.1111/1749-4877.12268 |
| **Various amphibian** | Body mass | Stark, G, Pincheira‐Donoso, D,  Meiri, S.  No evidence for the ‘rate‐of‐living’ theory across the tetrapod tree of life.  Global Ecol Biogeogr.  2020; 00:  1– 28. https: //doi.org/10.1111/geb.13069 |
| **Various amphibian** | SVL | AmphibiaWeb. 2020. <https://amphibiaweb.org> University of California, Berkeley, CA, USA. |
| **Various amphibians** | Body Mass | Trochet A, Moulherat S, Calvez O, Stevens V, Clobert J, Schmeller D (2014) A database of life-history traits of European amphibians. Biodiversity Data Journal 2: e4123. |
| **Various avian** | Body Mass | Terje Lislevand, Jordi Figuerola, and Tamás Székely. 2007. Avian body sizes in relation to fecundity, mating system, display behavior, and resource sharing. Ecology 88: 1605 |
| **Various avian** | Body Mass | Renner, S.C.; Hoesel, W. Ecological and Functional Traits in 99 Bird Species over a Large-Scale Gradient in Germany. Data 2017, 2, 12. |
| **Various mammals** | Body Mass | Smith, F.A., Lyons, S.K., Ernest, S.K.M., Jones, K.E., Kaufman, D.M., Dayan, T., Marquet, P.A., Brown, J.H. and Haskell, J.P. (2003), Body mass of late quaternary mammals. Ecology, 84: 3403-3403. |
| **Various primates** | Body Mass | Galán-Acedo, C., Arroyo-Rodríguez, V., Andresen, E. et al. Ecological traits of the world’s primates. Sci Data 6, 55 (2019) doi: 10.1038/s41597-019-0059-9 |
| **Various vertebrates** | Body Mass | Anthony I. Dell, Samraat Pawar, Van M. Savage. 2013. The thermal dependence of biological traits. Ecology 94: 1205. |

**Table S2 | Terrestrial model coefficients**

Estimates for all terrestrial classes, including those for splined latitude variable. Estimates in **bold** denote those included and fully discussed within the main text. Other estimates are included here, but not within the main text, due to their being uninterpretable.

| *Terrestrial* | *Estimate (ß)* | *SE* | *Lower CI* | *Upper CI* | *p-value* |
| --- | --- | --- | --- | --- | --- |
| *Birds* | | | | | |
| **(Intercept)** | **-1.455** | **0.261** | **-1.968** | **-0.943** | **0.082** |
| **Body Mass** | **-0.834** | **0.048** | **-0.928** | **-0.741** | **0.001** |
| Latitude (1) | -0.226 | 0.308 | -0.830 | 0.378 | 0.012 |
| Latitude (2) | -2.370 | 0.317 | -2.991 | -1.749 | 0.000 |
| Latitude (3) | 1.743 | 0.661 | 0.447 | 3.039 | 0.000 |
| Latitude (4) | -0.190 | 0.522 | -1.213 | 0.832 | 0.121 |
| Body Mass: Latitude (1) | -1.090 | 0.054 | -1.196 | -0.984 | 0.097 |
| Body Mass: Latitude (2) | -0.865 | 0.051 | -0.964 | -0.766 | 0.007 |
| Body Mass: Latitude (3) | -1.427 | 0.119 | -1.661 | -1.194 | 0.000 |
| Body Mass: Latitude (4) | -1.074 | 0.079 | -1.229 | -0.920 | 0.347 |
| *Mammals* | | | | | |
| **(Intercept)** | **-1.317** | **0.145** | **-1.601** | **-1.033** | **0.029** |
| **Body Mass** | **-0.952** | **0.008** | **-0.967** | **-0.936** | **0.000** |
| Latitude (1) | -0.164 | 0.117 | -0.392 | 0.065 | 0.000 |
| Latitude (2) | -1.026 | 0.116 | -1.254 | -0.799 | 0.821 |
| Latitude (3) | -0.383 | 0.295 | -0.961 | 0.196 | 0.037 |
| Latitude (4) | -1.157 | 0.130 | -1.411 | -0.902 | 0.228 |
| *Amphibians* | | | | | |
| **(Intercept)** | **0.074** | **0.193** | **-0.304** | **0.453** | **0.000** |
| **Body Mass** | **-1.172** | **0.078** | **-1.325** | **-1.018** | **0.028** |
| *Reptiles* | | | | | |
| **(Intercept)** | **-0.118** | **0.248** | **-0.603** | **0.368** | **0.000** |
| **Body Mass** | **-0.987** | **0.059** | **-1.103** | **-0.872** | **0.832** |
| Latitude (1) | -1.703 | 0.381 | -2.449 | -0.956 | 0.065 |
| Latitude (2) | -1.047 | 0.295 | -1.625 | -0.469 | 0.874 |
| Latitude (3) | -1.325 | 0.759 | -2.813 | 0.163 | 0.668 |
| Latitude (4) | -1.949 | 0.586 | -3.098 | -0.801 | 0.105 |
| Body mass: Latitude (1) | -0.901 | 0.068 | -1.033 | -0.768 | 0.142 |
| Body mass: Latitude (2) | -0.967 | 0.052 | -1.068 | -0.865 | 0.522 |
| Body mass: Latitude (3) | -1.030 | 0.154 | -1.331 | -0.729 | 0.845 |
| Body mass: Latitude (4) | -0.884 | 0.119 | -1.118 | -0.650 | 0.330 |

**Table S3 | Marine model coefficients**

Estimates for all marine classes, including those for splined latitude variable. Estimates in **bold** denote those included and fully discussed within the main text. Other estimates are included here, but not within the main text, due to their being uninterpretable.

| *Marine* | *Estimate (ß)* | *SE* | *Lower CI* | *Upper CI* | *p-value* |
| --- | --- | --- | --- | --- | --- |
| *Birds* | | | | | |
| **(Intercept)** | **-3.804** | **1.967** | **-7.659** | **0.051** | **0.154** |
| **Body Mass** | **-0.710** | **0.228** | **-1.156** | **-0.263** | **0.203** |
| Latitude (1) | 1.382 | 1.974 | -2.486 | 5.250 | 0.227 |
| Latitude (2) | 3.212 | 1.583 | 0.110 | 6.314 | 0.008 |
| Latitude (3) | 4.015 | 3.771 | -3.375 | 11.406 | 0.184 |
| Latitude (4) | -0.068 | 0.970 | -1.970 | 1.834 | 0.337 |
| Body Mass: Latitude (1) | -1.122 | 0.235 | -1.582 | -0.662 | 0.604 |
| Body Mass: Latitude (2) | -1.469 | 0.191 | -1.842 | -1.095 | 0.014 |
| Body Mass: Latitude (3) | -1.316 | 0.439 | -2.177 | -0.455 | 0.472 |
| Body Mass: Latitude (4) | -1.050 | 0.129 | -1.303 | -0.797 | 0.698 |
| *Mammals* | | | | | |
| **(Intercept)** | **-0.098** | **0.331** | **-0.747** | **0.550** | **0.006** |
| **Body Mass** | **-1.018** | **0.025** | **-1.067** | **-0.969** | **0.467** |
| *Reptiles* | | | | | |
| **(Intercept)** | **0.117** | **0.306** | **-0.484** | **0.717** | **0.000** |
| **Body Mass** | **-1.017** | **0.027** | **-1.069** | **-0.964** | **0.529** |
| *Bony* *Fish* |  |  |  |  |  |
| **(Intercept)** | **0.365** | **0.591** | **-0.794** | **1.524** | **0.021** |
| **Body Mass** | **-1.101** | **0.066** | **-1.231** | **-0.971** | **0.128** |
| Latitude (1) | -1.127 | 0.540 | -2.185 | -0.068 | 0.814 |
| Latitude (2) | -2.047 | 0.458 | -2.945 | -1.149 | 0.022 |
| Latitude (3) | -3.867 | 1.264 | -6.344 | -1.390 | 0.023 |
| Latitude (4) | -1.484 | 0.315 | -2.102 | -0.866 | 0.125 |
| Body Mass: Latitude (1) | -0.966 | 0.062 | -1.087 | -0.845 | 0.583 |
| Body Mass: Latitude (2) | -0.896 | 0.053 | -0.999 | -0.792 | 0.048 |
| Body Mass: Latitude (3) | -0.673 | 0.143 | -0.953 | -0.393 | 0.022 |
| Body Mass: Latitude (4) | -0.963 | 0.040 | -1.041 | -0.886 | 0.357 |
| *Cartilaginous Fish* | | | | | |
| **(Intercept)** | **-0.281** | **0.116** | **-0.508** | **-0.053** | **0.000** |
| **Body Mass** | **-0.966** | **0.010** | **-0.986** | **-0.946** | **0.001** |
| Latitude (1) | -1.315 | 0.114 | -1.539 | -1.091 | 0.006 |
| Latitude (2) | -1.253 | 0.079 | -1.408 | -1.098 | 0.001 |
| Latitude (3) | -2.132 | 0.216 | -2.555 | -1.708 | 0.000 |
| Latitude (4) | -1.327 | 0.124 | -1.570 | -1.085 | 0.008 |

**Table S4 | Freshwater model coefficients**

Estimates for all freshwater classes, including those for splined latitude variable. Estimates in **bold** denote those included and fully discussed within the main text. Other estimates are included here, but not within the main text, due to their being uninterpretable.

| *Freshwater* | *Estimate (ß)* | *SE* | *Lower CI* | *Upper CI* | *p-value* |
| --- | --- | --- | --- | --- | --- |
| *Birds* | | | | | |
| **(Intercept)** | **-0.556** | **0.184** | **-0.917** | **-0.195** | **0.016** |
| **Body Mass** | **-1.016** | **0.018** | **-1.051** | **-0.980** | **0.398** |
| Latitude (1) | -0.364 | 0.144 | -0.646 | -0.082 | 0.000 |
| Latitude (2) | -1.220 | 0.120 | -1.455 | -0.985 | 0.066 |
| Latitude (3) | -0.099 | 0.441 | -0.963 | 0.765 | 0.041 |
| Latitude (4) | -0.411 | 0.186 | -0.775 | -0.046 | 0.002 |
| *Mammals* | | | | | |
| **(Intercept)** | **-2.245** | **0.592** | **-3.406** | **-1.084** | **0.036** |
| **Body Mass** | **-0.895** | **0.042** | **-0.977** | **-0.813** | **0.012** |
| Latitude (1) | 0.047 | 0.297 | -0.535 | 0.629 | 0.000 |
| Latitude (2) | -1.088 | 0.361 | -1.796 | -0.381 | 0.807 |
| Latitude (3) | 1.608 | 0.778 | 0.082 | 3.133 | 0.001 |
| Latitude (4) | -0.600 | 0.249 | -1.088 | -0.111 | 0.108 |
| *Bony* *Fish* | | | | | |
| **(Intercept)** | **-0.281** | **0.306** | **-0.882** | **0.320** | **0.019** |
| **Body Mass** | **-1.062** | **0.042** | **-1.145** | **-0.979** | **0.142** |
| Latitude (1) | -0.079 | 0.366 | -0.796 | 0.638 | 0.012 |
| Latitude (2) | -1.811 | 0.275 | -2.350 | -1.272 | 0.003 |
| Latitude (3) | -1.279 | 0.801 | -2.849 | 0.291 | 0.728 |
| Latitude (4) | 1.336 | 0.459 | 0.437 | 2.235 | 0.000 |
| Body Mass: Latitude (1) | -1.036 | 0.047 | -1.128 | -0.943 | 0.450 |
| Body Mass: Latitude (2) | -0.893 | 0.034 | -0.959 | -0.828 | 0.001 |
| Body Mass: Latitude (3) | -0.851 | 0.110 | -1.067 | -0.636 | 0.176 |
| Body Mass: Latitude (4) | -1.307 | 0.051 | -1.407 | -1.206 | 0.000 |
| *Amphibians* | | | | | |
| **(Intercept)** | **-2.712** | **1.249** | **-5.160** | **-0.265** | **0.170** |
| **Body Mass** | **-0.060** | **0.441** | **-0.925** | **0.805** | **0.033** |
| Latitude (1) | -1.029 | 1.187 | -3.355 | 1.297 | 0.981 |
| Latitude (2) | 3.641 | 0.944 | 1.791 | 5.492 | 0.000 |
| Latitude (3) | 2.742 | 3.341 | -3.806 | 9.290 | 0.263 |
| Latitude (4) | -12.558 | 3.463 | -19.345 | -5.770 | 0.001 |
| Body Mass: Latitude (1) | -1.337 | 0.417 | -2.155 | -0.519 | 0.419 |
| Body Mass: Latitude (2) | -2.148 | 0.274 | -2.686 | -1.610 | 0.000 |
| Body Mass: Latitude (3) | -3.380 | 1.148 | -5.629 | -1.130 | 0.038 |
| Body Mass: Latitude (4) | 1.616 | 0.747 | 0.151 | 3.080 | 0.000 |
| *Reptiles* | | | | | |
| **(Intercept)** | **0.200** | **0.256** | **-0.303** | **0.702** | **0.000** |
| **Body mass** | **-0.988** | **0.017** | **-1.021** | **-0.954** | **0.460** |
| Latitude (1) | -1.434 | 0.241 | -1.905 | -0.962 | 0.072 |
| Latitude (2) | -0.935 | 0.203 | -1.334 | -0.537 | 0.751 |
| Latitude (3) | -2.082 | 0.492 | -3.047 | -1.117 | 0.028 |
| Latitude (4) | -0.752 | 0.170 | -1.085 | -0.419 | 0.145 |
